# Molecular phylogenetics supports a clade of red algal parasites retaining native plastids: taxonomy and terminology revised

**DOI:** 10.1101/182709

**Authors:** Eric D. Salomaki, Christopher E. Lane

## Abstract

Parasitism is a life strategy that has repeatedly evolved within the Florideophyceae. Historically, the terms adelphoparasite and alloparasite have been used to distinguish parasites based on the relative phylogenetic relationship of host and parasite. However, analyses using molecular phylogenetics indicate that nearly all red algal parasites infect within their taxonomic family, and a range of relationships exist between host and parasite. To date, all investigated adelphoparasites have lost their plastid, and instead, incorporate a host derived plastid when packaging spores. In contrast, a highly reduced plastid lacking photosynthesis genes was sequenced from the alloparasite *Choreocolax polysiphoniae*. Here we present the complete *Harveyella mirabilis* plastid genome, which has also lost genes involved in photosynthesis, and a partial plastid genome from *Leachiella pacifica*. The *H. mirabilis* plastid shares more synteny with free-living red algal plastids than that of *C. polysiphoniae*. Phylogenetic analysis demonstrates that *C. polysiphoniae, H. mirabilis*, and *L. pacifica* form a robustly supported clade of parasites, which retain their own plastid genomes, within the Rhodomelaceae. We therefore transfer all three genera from the exclusively parasitic family, Choreocolacaceae, to the Rhodomelaceae. Additionally, we recommend applying the terms archaeplastic parasites (formerly alloparasites), and neoplastic parasites (formerly adelphoparasites) to distinguish red algal parasites using a biological framework rather than taxonomic affiliation with their hosts.

## INTRODUCTION

Since the late 19^th^ century when parasitic red algae were first formally recognized the number of red algal parasite species has been steadily increasing (Reinsch 1875, Setchell 1914, 1923, Goff 1982, Preuss et al. 2017). Recent counts identify 124 distinct species of red algal parasites distributed across eight of the 31 of the Florideophyceae (Salomaki and Lane 2014, Blouin and Lane 2015, Preuss et al. 2017, Preuss and Zuccarello 2018, Guiry and Guiry 2018). The abundance of successful independent evolutions of parasitism within the Florideophyceae seems in large part due to their ability to form direct cell-to-cell fusions between adjacent, non-daughter cells (Wetherbee and Quirk 1982a, Goff and Coleman 1985). With few exceptions [e.g. *Choreonema thuretii* (Broadwater and LaPointe 1997)], red algal parasites use their ability to form cell fusions as a means of infecting their host (Goff and Coleman 1985, Zuccarello et al. 2004).

Red algal parasites exclusively infect other rhodophytes and are predominately unpigmented, appearing as galls or irregular growths on their free-living red algal hosts. Despite their diminutive habit, parasitic red algae share morphological characteristics with other close relatives, and thus were assigned to tribes or families at the time of their initial discovery (Reinsch 1875, Feldmann and Feldmann 1958). In the early 20^th^ century, two hypotheses arose to explain the origins of red algal parasites. First, Setchell (1918) noted that approximately 80% of the parasites he was studying infected hosts in the same family. Based upon his observations, he proposed that parasites originated as carpospores or tetraspores of their host, which had undergone a mutation causing reduced photosynthetic capabilities (Setchell 1918). Later, Sturch (1926) proposed that rather than evolving sympatrically, parasitic red algae started out as small epiphytes of their hosts that eventually would penetrate cortical cells of the host, becoming an endophyte. Once established as an endophyte, the alga would adopt mechanisms to obtain nutrients from the host, becoming increasingly reliant on its host, solidifying an irreversible path towards parasitism (Sturch 1926). Feldmann and Feldmann (1958) furthered Sturch’s hypothesis, adding that epiphytes that are closely related to their hosts are more likely to succeed in creating connections with hosts cells and are therefore more likely to complete a transition to parasitism.

Historically parasites that have infected species within their own family or tribe have been considered ‘adelphoparasites’, while those more distantly related to their hosts are called ‘alloparasites’ (Feldmann and Feldmann 1958, Goff et al. 1996). In agreement with Setchell’s initial observations (1918), approximately 80% of the currently described red algal parasite species are considered to be adelphoparasitic, while the remaining 20% are alloparasitic (Goff 1982, Salomaki and Lane 2014, Blouin and Lane 2015). Based initially on morphological observations, and later coupled with molecular data, it was proposed that parasites evolve sympatrically with their hosts, as adelphoparasites, and over time diversify or adapt to infect new and more distantly related hosts, becoming alloparasites (Feldmann and Feldmann 1958, Blouin and Lane 2012).

Sturch (1926) described the Choreocolacaceae as a family in the Gigartinales comprised of morphologically reduced parasites lacking chlorophyll including species of *Choreocolax, Harveyella*, and *Holmsella*. This exclusively parasitic family was the subject of a thorough phylogenetic analysis of alloparasites to confirm whether these parasitic red algae were truly monophyletic (Zuccarello et al. 2004). Their study supported previous morphological observations (Fredericq and Hommersand 1990) confirming that *Holmsella* was a member of the Gracilariaceae and questioned the legitimacy of recognizing a family of red algal parasites. However, Zuccarello et al. (2004) did not formally address the taxonomic implications for the Choreocolacaceae.

In the few other cases where molecular phylogenetics have been applied to assess evolutionary histories of red algal parasites, data suggests that red algal parasites have arisen though numerous independent evolutionary events (Goff et al. 1996, 1997, Kurihara et al. 2010). In addition to phylogenetic analyses, molecular tools have also been applied to investigate the parasite-host dynamics throughout parasite development. Analyses of the adelphoparasites *Gardneriella tuberifera* Kylin, *Gracilariophila oryzoides* Setchell & H. L. Wilson demonstrated that, although the parasites maintain a native mitochondrion, they have lost their native plastid and instead ‘hijack’ a host plastid when packaging their own spores (Goff and Coleman 1995). To date, all red algal parasites examined maintain a fully functional mitochondrion (Salomaki and Lane 2016). All adelphoparasites that have been investigated have lost their native plastid (Goff and Coleman 1995; Salomaki and Lane 2014), and next-generation sequencing efforts targeting *Faucheocolax attenuatus* Setchell, *Janczewskia gardneri* Setchell & Guernsey, and *G. oryzoides* support these findings (Salomaki and Lane unpublished). In contrast, a highly reduced native plastid was sequenced from the alloparasite *Choreocolax polysiphoniae* Reinsch, which lost genes involved in photosynthesis, yet maintains functions including fatty acid and amino acid biosynthesis (Salomaki et al. 2015). The lack of plastids in adelphoparasites, in combination with finding a native plastid in the alloparasite *C. polysiphoniae*, demonstrates that not all parasites pass through an ‘adelphoparasite stage’ and that there are multiple paths to parasitism in red algae.

In their study examining relationships in the Choreocolacaceae, Zuccarello *et al*. (2004) found that *Holmsella pachyderma* and *Holmsella australis* formed a clade within the Gracilariaceae. Additionally they demonstrated that the parasite genera *Choreocolax*, *Harveyella*, and *Leachiella* belonged in the Rhodomelaceae, but lacked support for a monophyletic group of parasites. Using DNA sequence data we investigated the relationships of *Choreocolax, Harveyella*, and *Leachiella*, and tested the hypothesis that parasites can arise and subsequently speciate, forming clades that infect a range of hosts. Furthermore, we explored the evolution of parasitism and the loss of photosynthesis among species traditionally called alloparasites.

## MATERIALS AND METHODS

### Sample Collection and DNA Extraction

*Harveyella mirabilis* (Reinsch) F.Schmitz & Reinke, on its host, *Odonthalia washingtoniensis* Kylin, was collected from Cattle Point, Friday Harbor, WA, USA (48.452428, −122.962774), and *Leachiella pacifica* Kugrens, on *Polysiphonia hendryi* N.L. Gardner, was collected off the dock at Friday Harbor Laboratories, Friday Harbor WA, USA (48.545066, −123.012268). Individual parasite galls from *Harveyella mirabilis* and *Leachiella pacifica* were dissected from their respective hosts and collected in a 1.5 mL microcentrifuge tube. Unfortunately, only single parasites of *H. mirabilis* and *L. pacifica* were observed on the host tissue and the entire pustule was excised for DNA isolation. Dried vouchers for the remaining host tissue from *Polysiphonia hendryi* (specimen 03304646) and *Odonthalia washingtoniensis* (specimen 03304647) have been deposited at the New York Botanical Garden herbarium (NY). Parasite tissue was hand-ground using a Corning Axygen^®^ PES-15-B-SI disposable tissue grinder pestle in a 1.5 mL microcentrifuge tube while submerged in 100μL of DNA extraction buffer (Saunders 1993). DNA was extracted from specimens using a standard phenol/chloroform extraction with all ratios adjusted for an initial buffer volume of 100μL (Saunders 1993).

### Molecular Analyses

Genomic DNA was amplified from *Harveyella mirabilis* and *Leachiella pacifica* using the illustra Single Cell GenomiPhi DNA Amplification kit (GE Healthcare Life Sciences, Pittsburgh, Pa) according to manufacturer protocols. Libraries for Illumina sequencing were constructed from the amplified DNA on the Apollo 324 robot using the PrepX ILM DNA Library Kit (Wafergen Biosystems, Freemont, CA, USA). The *Harveyella mirabilis* and *Leachiella pacifica* libraries were multiplexed and sequenced on an Illumina MiSeq paired-end 250 x 250 basepair run. The sequencing effort resulted in 7,923,094 paired end reads for *Harveyella mirabilis* and 7,502,360 paired-end reads for *Leachiella pacifica*. Raw sequencing reads with Phred scores <30 were removed and the remaining reads were trimmed of adapter sequences. Additionally, fifteen 5’ and five 3’ nucleotides were trimmed from the remaining reads and all reads under 100 nucleotides were removed from the dataset. All trimming was completed using CLC Genomics Workbench v. 9.5.2 (CLC Bio-Qiagen, Aarhus, Denmark) and the remaining reads were assembled using default parameters in CLC Genomics Workbench v. 9.5.2.

### Plastid Genome Annotation

A 90,654 bp contig was identified as the plastid genome of *Harveyella mirabilis* from the assembled Illumina MiSeq data (GenBank Accession MK039118). We recovered >40,000 bp of the *Leachiella pacifica* plastid genome that were in five separate contigs (GenBank Accession MK039114-MK039117, MK039119). Open reading frame (ORF) prediction on the plastid genomes was done in Geneious Pro v. 9.1.5 and the resulting ORFs were manually annotated using GenBank and Pfam (Finn et al. 2010, 2015) databases. Functional annotations were assigned from the UniProt (The UniProt Consortium 2017) and KEGG databases (Kanehisa et al. 2016). Genes found in red algal plastid genomes that were missing from the *H. mirabilis* plastid were searched for using BLAST, against the plastid sequence and the genomic assemblies to verify their absence and check for evidence of transfer to another genetic compartment. Plastid genome sequences were submitted to the tRNAscan-SE online server v1.21 (Schattner et al. 2005) for identification of tRNA sequences and to MFannot (http://megasun.bch.umontreal.ca/cgi-bin/mfannot/mfannotInterface.pl) to identify rRNA sequences and confirm manual gene annotations.

### Plastid Genome Comparative Analysis

The circular plastid genomes of from the florideophytes, *Calliarthron tuberculosum* (KC153978), *Ceramium cimbricum* (KR025491), *Ceramium japonicum* (KX284719), *Chondrus crispus* (HF562234), *Dasya binghamiae* (KX247284), *Laurencia sp*. JFC0032 (LN833431), *Grateloupia taiwanensis* (KC894740), *Gracilaria tenuistipitata* (AY673996), *Vertebrata lanosa* (KP308097), and the alloparasites *Choreocolax polysiphoniae* (KP308096) and *Harveyella mirabilis* (MK039118) were arranged so that their sequences began with the *fts*H gene. Whole genome alignment was completed using the default settings for the progessiveMauve algorithm in the Mauve v2.3.1 (Darling et al. 2004).

### Phylogenetic Analysis

A data matrix comprising 17 protein coding genes (*acc*B, *acp*P, *fab*H, *ilv*H, *odp*A, *rne*, *rpl*16, *rpl*20, *rpl*3, *rps*14, *rps*16, *rps*19, *suf*B, *trp*A, *trp*G, *tsf ycf*19) and two plastid encoded rRNAs (*rnl*, *rns*) that were shared between the *Harveyella mirabilis* plastid, the partial *Leachiella pacifica* plastid, and 55 completely sequenced plastid genomes from the Ceramiales was assembled to investigate the placement of *Harveyella mirabilis* and *Leachiella pacifica*, within the Rhodomelaceae (Salomaki et al. 2015, Verbruggen and Costa 2015, Hughey and Boo 2016, Díaz-Tapia et al. 2017). Shared plastid genes were extracted into 19 distinct datasets in Geneious Pro v. 9.1.5 (Kearse et al. 2012). The protein coding gene datasets were independently aligned using the translation align setting in Geneious v. 9.1.5 and the rRNAs were aligned using the MAFFT online server (Katoh et al. 2017) and subsequently concatenated using Sequence Matrix v. 1.7.8 (Vaidya et al. 2011) producing a 15,763 site dataset. The best-fit partitioning scheme and the best model of evolution for each partition was inferred using the Bayesian information criterion (BIC) as implemented in PartitionFinder2 (Lanfear et al. 2017). The concatenated dataset was then subjected to phylogenetic analysis using maximum likelihood, implemented in both RAxML v. 8.2.2 (Stamatakis 2014) and IQ-TREE v. 1.5.6 (Nguyen et al. 2015), as well as Bayesian Inference using MrBayes v. 3.2.2 (Ronquist et al. 2012). For the maximum likelihood analysis the bootstrap support values were calculated using 1,000 bootstrap replicates in RAxML, and 1,000 replicates of ultrafast bootstrapping (UFBoot) in the IQ-TREE analysis (Hoang et al. 2018). For the Bayesian inference analysis, two Metropolis-coupled Markov chain Monte Carlo (MCMCMC) runs consisting of one cold chain and three hot chains were preformed. Each run was sampled every 500 generations for 2,000,000 generations. After confirming the runs converged by checking to ensure that the average standard deviation of split frequencies was below 0.01, the trees were merged. The resulting tree and posterior probabilities were calculated from the 8,002 trees generated.

## RESULTS

### An Exclusively Parasitic Clade

Phylogenetic analysis was conducted on a concatenated dataset comprised of 19 plastid-encoded genes from 54 free-living species, and the parasites *C. polysiphoniae, H. mirabilis, L. pacifica*. A Bayesian inference analysis implemented in MrBayes and maximum likelihood analyses using both RAxML and IQ-TREE all recovered robust support for a topology similar to that recovered by Díaz-Tapia et al. (2017). The three parasites formed a clade with 1.0 posterior probability (BI), 100% bootstrap support in RAxML and 100% ultrafast bootstrap approximation (UFBoot) in IQ-TREE (Fig. 1). *Choreocolax polysiphoniae* was the earliest branching parasite with *H. mirabilis* and *L. pacifica* being recovered as sister taxa with 100% bootstrap support/1.0 posterior probability (Fig. 1).

**Figure 1.**
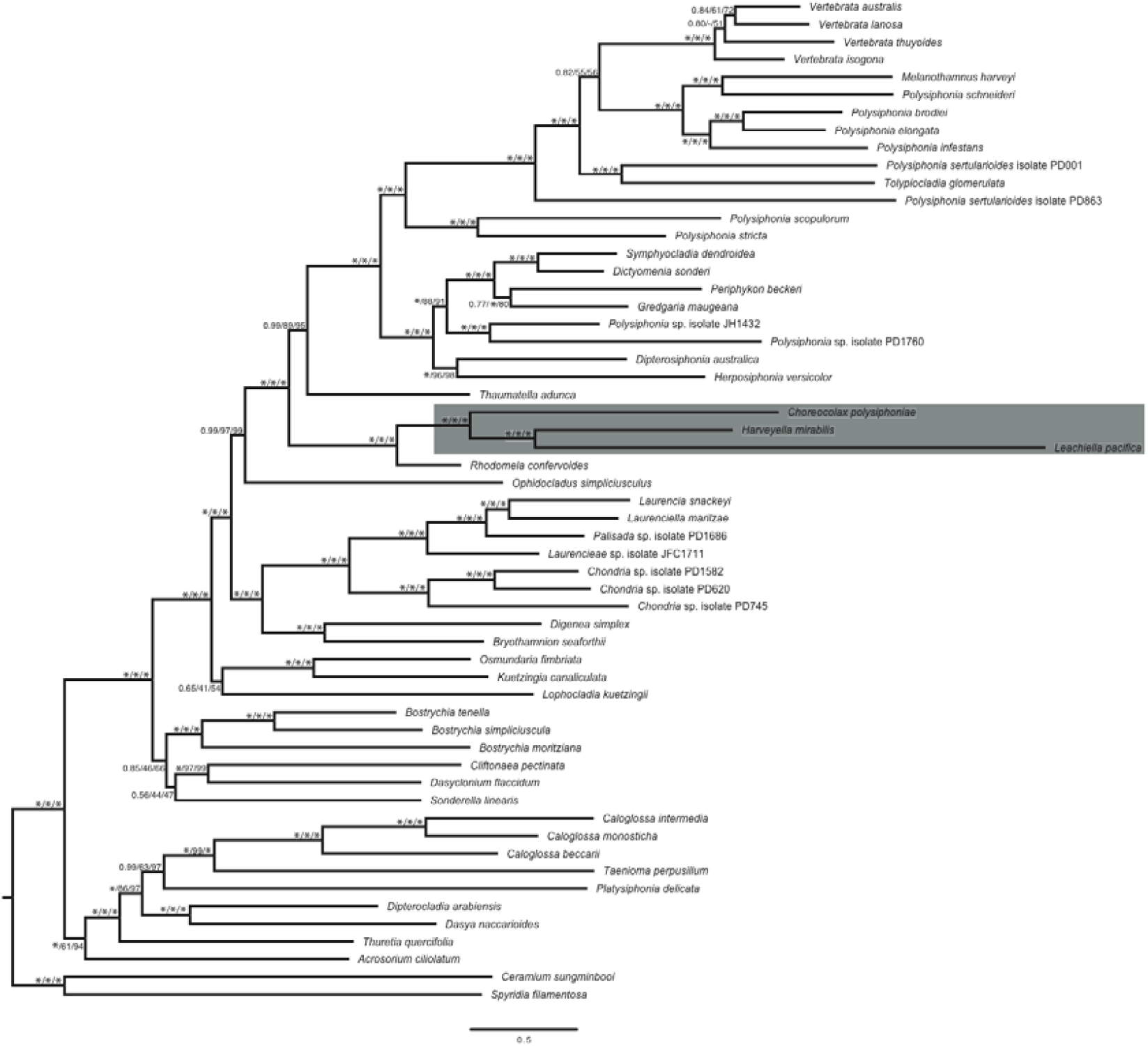
Maximum likelihood phylogeny of the parasites *Choreocolax polysiphoniae, Harveyella mirabilis, Leachiella pacifica*, and 54 free-living members of the Ceramiales. This analysis provides full support for a clade of parasites arising within the Rhodomelaceae. Support values shown as Bayesian posterior probability/RAxML bootstrap/IQ-TREE UFboot.

### Taxonomic Considerations

*Choreocolax polysiphoniae* was initially described by Reinsch (1875) from the Atlantic coast of North America as parasitic on *Polysiphonia fastigiata* (now *Vertebrata lanosa*). Specimens used in the phylogenetic analysis presented here (Fig. 1) were collected at Beavertail State Park in Jamestown, RI, USA infecting *Vertebrata lanosa* and are a strong match to the type description (Reinsch 1875). A representative voucher of *C. polysiphoniae* on its host *V. lanosa* (03304648) has been deposited at the New York Botanical Garden herbarium (NY). As the type collection cannot be located, we formally lectotypify *C. polysiphoniae* on image #49 accompanying the description in Reinsch (1875). Based upon the molecular analyses here, which resolved a monophyletic clade of parasitic red algae within the Rhodomelaceae (Fig. 1), we formally transfer *Choreocolax* to the Rhodomelaceae and recognize Choreocolacaceae as a synonym of this family. To adhere to the principle of monophyly, the genus *Harveyella*, based on the type and only species *H. mirabilis* and included in our analyses (Fig. 1), is also transferred to the Rhodomelaceae.

### Parasite Plastid Genomes

The complete plastid genome of *Harveyella mirabilis* was assembled as a 90,654 kb circular molecule with 322x coverage. The plastid genome has an overall AT content of 76.5% and contains 84 protein coding genes, 3 rRNAs, and 23 tRNAs (Fig. 2). Similar to the *Choreocolax polysiphoniae* plastid (Salomaki et al. 2015), all genes related to photosynthesis have been lost with the exception of petF, which is involved in electron transport in other metabolic pathways (Happe and Naber 1993, Jacobs et al. 2009). Genes involved in transcription/translation and fatty acid, amino acid, protein, isoprene biosynthesis remain conserved. As in the *C. polysiphoniae* plastid, *glt*B appears to be a pseudogene. BLAST similarity searches are able to find conserved homology, however the presence of stop-codons throughout the region suggests that the gene is likely no longer capable of being completely translated.

**Figure 2.**
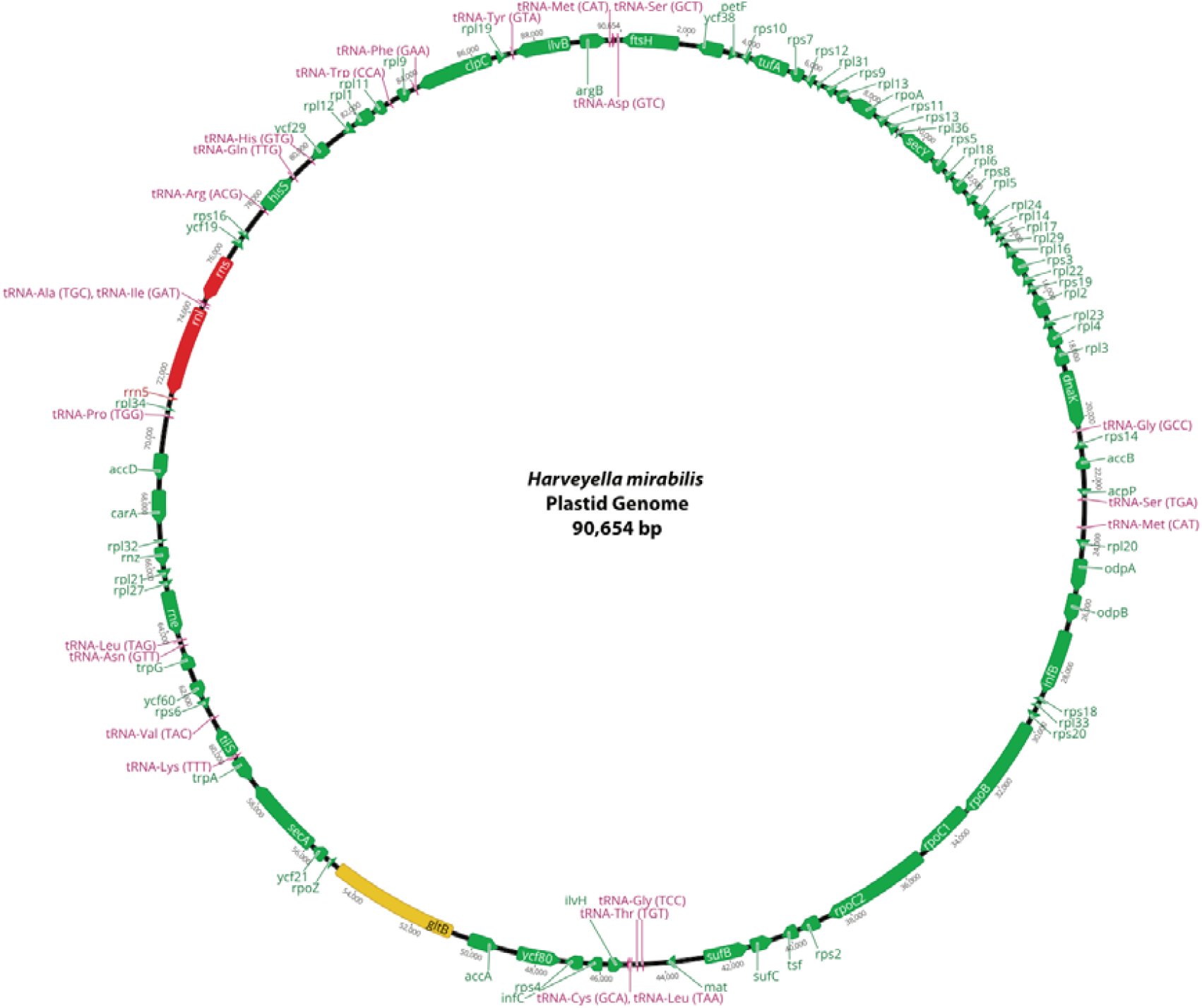
The plastid genome of the parasitic red alga, *Harveyella mirabilis* is 90,654 basepairs and contains 84 protein-coding genes (Green), the 5S, 16S, and 23S rRNAs (Red), and 23 tRNAs (Pink). All genes involved with photosynthetic functions, except *pet*F, have been lost. The *fts*H gene is truncated but may still be transcribed, however *glt*B is a non-functional pseudogene (Yellow).

Five contigs, comprising 40,895 bp of the *Leachiella pacifica* plastid genome, were assembled and annotated. These genomic pieces encode 24 complete protein-coding genes, 6 protein-coding gene fragments at the ends of contigs, the small and large rRNA subunits, and 12 tRNAs. Similar to *H. mirabilis* and *C. polysiphoniae*, no genes associated with photosynthesis were recovered from the incomplete *L. pacifica* plastid genome.

### Plastid Genome Comparisons

A whole genome MAUVE alignment of nine representative free-living Florideophyceae plastid genomes, the *H. mirabilis* and *C. polysiphoniae* genomes, identified 13 locally collinear blocks in the *H. mirabilis* genome that aligned with the free-living plastids (Fig. 3). There were no rearrangements or inversions in the *H. mirabilis* plastid genome when compared to photosynthetic Rhodomelaceae taxa. When aligning *H. mirabilis* to the *C. polysiphoniae* plastid genome, 11 locally collinear blocks were identified, and several genome inversions and rearrangements are evident (Fig. 4). Additionally, gene content varies slightly between the two parasite plastid genomes. *Harveyella mirabilis* retains *arg*B, *car*A, *inf*B, *rnz*, *rpl*9, *rpl*17, *rpl*24, *rpl*32, *rpl*33, *rpl*34, *rpo*Z, *rps*18, *rps*20, *ycf*21, which have all been lost in *C. polysiphoniae* (Table 1), while *C. polysiphoniae* maintains copies of *dna*B and *fab*H which have both been lost in the *H. mirabilis* plastid genome.

**Figure 3.**
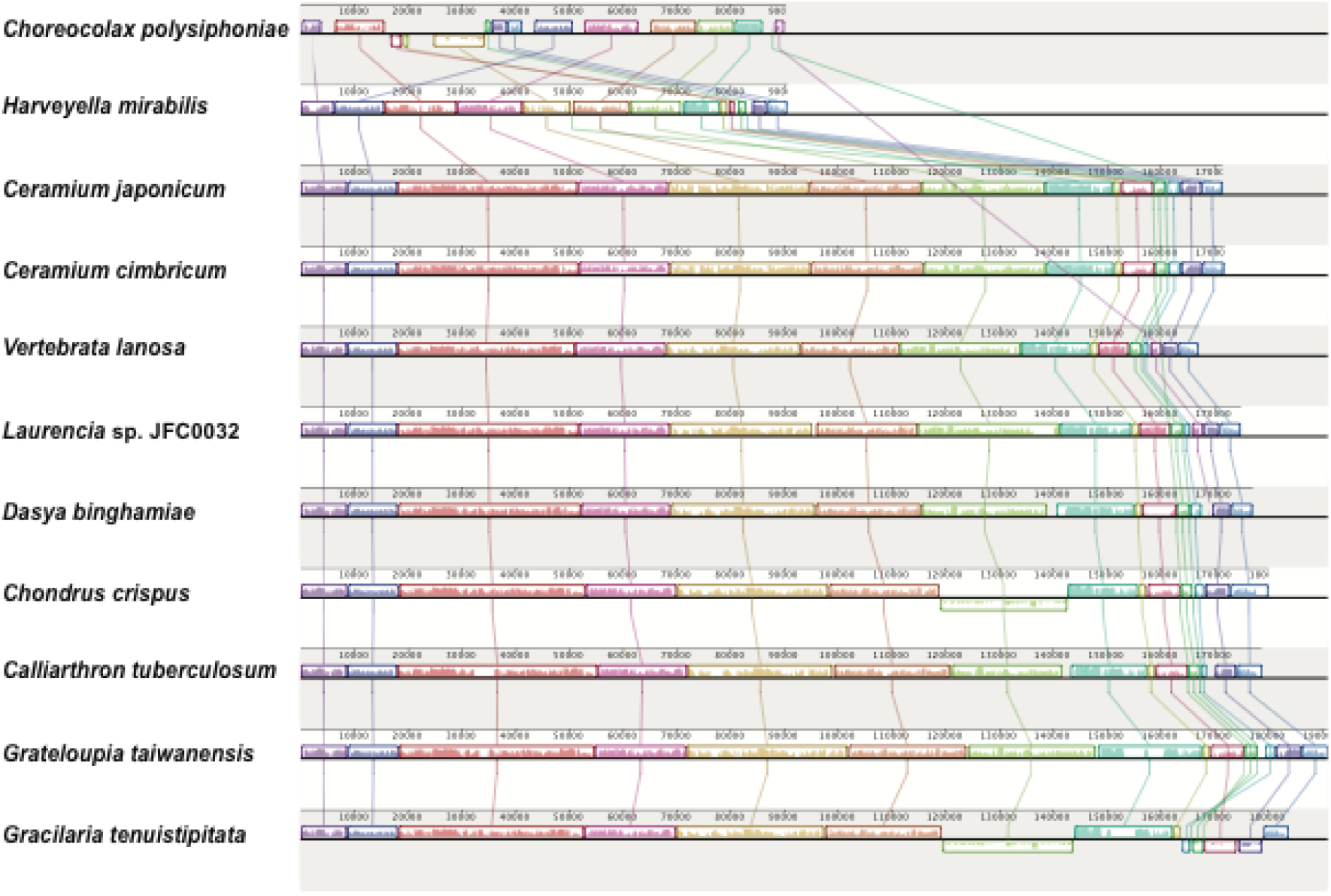
Mauve alignment of the parasites *Choreocolax polysiphoniae* (top) and *Harveyella mirabilis* (second from top) with all published Rhodomelaceae plastid genomes, as well as those from select representatives of other Florideophyceae families (*Dasya binghamiae*, Dasyaceae; *Chondrus crispus*, Gigartinaceae; *Calliarthron tuberculosum*, Corallinaceae; *Grateloupia taiwanensis*, Halymeniaceae; *Gracilaria tenuistipitata*, Gracilariaceae). This alignment identifies 16 locally collinear blocks (LCBs) among the selected plastid genomes and demonstrates that even with the loss of photosynthesis genes overall synteny is shared between the parasite *H. mirabilis* and other plastid genomes, while *C. polysiphoniae* has undergone several genome rearrangements.

**Table 1.**
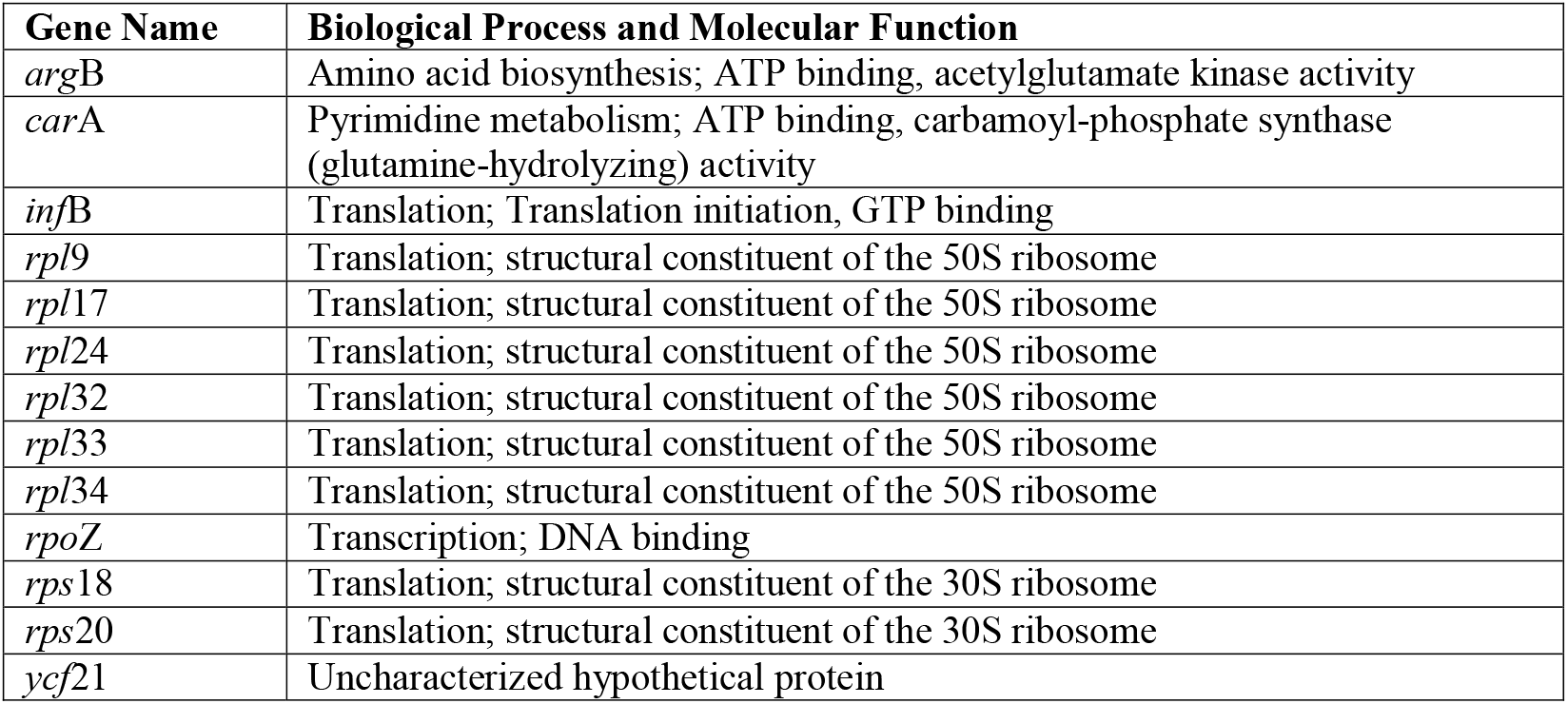
*Harveyella mirabilis* plastid genes and their function, which have been lost from the *Choreocolax polysiphoniae* plastid genome.

## DISCUSSION

### Parasite Plastids

The link between plastid origin in red algal parasites and their evolutionary relationship to their hosts may be central to our understanding of red algal parasite evolution. The *Harveyella mirabilis* plastid genome (Fig. 2) represents the second red algal parasite demonstrated to retain a reduced native plastid. Similar to *Choreocolax polysiphoniae*, the *H. mirabilis* plastid genome remains conserved for functions including amino acid, fatty acid, and protein biosynthesis, but has lost genes involved in building the light harvesting apparatus, photosystems I and II, and other photosynthesis related genes. The partial *Leachiella pacifica* plastid genome presented here also supports that hypothesis, although work remains to completely sequence its plastid genome.

The *H. mirabilis* plastid gene order, with the exception of the missing photosynthesis genes, is conserved when compared to plastids of free-living Rhodomelaceae species (Fig. 3). In contrast, the *C. polysiphoniae* plastid has undergone greater gene loss than the *H. mirabilis* plastid, and a substantial amount of genome reorganization (Fig. 4). *Harveyella mirabilis* retains 13 genes that have been lost from the *C. polysiphoniae* plastid (Table 1); *arg*B and *car*A, which are involved in arginine biosynthesis processes, *rpo*Z, which promotes RNA polymerase assembly, nine ribosomal protein genes, and an uncharacterized hypothetical protein. In contrast, *C. polysiphoniae* retains copies of *dna*B, which is involved in DNA replication, and *fab*H, which is involved in fatty acid biosynthesis, both of which have been lost in *H. mirabilis*. Analysis of additional plastid genomes in this clade will provide greater insights into patterns of plastid genome evolution in red algal parasites.

**Figure 4.**
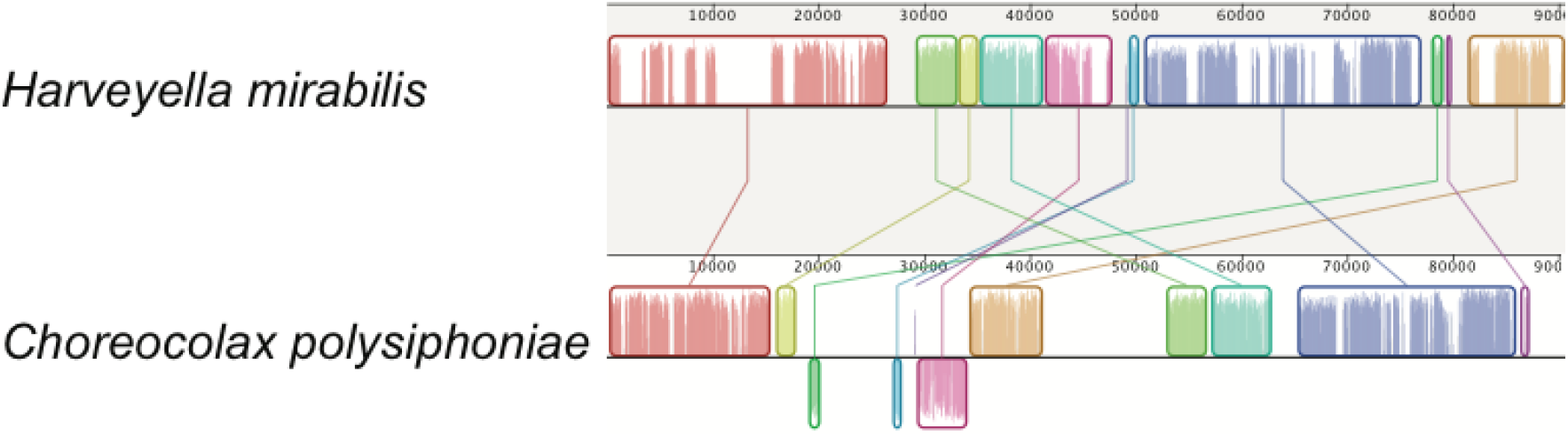
Mauve alignment of the plastid genomes from the parasites *Harveyella mirabilis* (top) and *Choreocolax polysiphoniae* (bottom). This alignment identifies 11 locally collinear blocks (LCBs) among these parasite plastid genomes, highlighting the high level of genome fragmentation and rearrangements evident in the two parasite plastids.

### The Demise of Exclusively Parasitic Families

Sturch (1926) initially described the Choreocolacaceae as a family of holoparasites, containing the genera *Choreocolax, Harveyella*, and *Holmsella*. However, more recent morphological investigation showed that *Holmsella* was related to the parasites *Gelidiocolax* and *Pterocladiophila*, and it was moved to the family Pterocladiophilaceae in the Gracilariales (Fredericq and Hommersand 1990). Their observations were subsequently supported by molecular data generated with the specific aim of testing the phylogenetic affinities of parasites that Sturch had assigned to the Choreocolacaceae. This work demonstrated that *Holmsella australis* and *Holmsella pachyderma* formed a well-supported monophyletic clade within the Gracilariaceae (Zuccarello et al. 2004). Their molecular data also indicated that *Choreocolax* and *Harveyella* belonged in the Ceramiales, leading the authors to question whether the Choreocolacaceae should continue to be recognized (Zuccarello et al. 2004). However, their use of 18S sequence data was insufficient to resolve a monophyletic clade exclusively comprised of parasites. Other authors have also noted that species in *Choreocolax, Harveyella*, and *Leachiella* have features aligning them to the Ceramiales, but again, taxonomic affinities among the parasite species and within this order remained uncertain (Goff and Cole 1975, Kugrens 1982, Fredericq and Hommersand 1990). Our data confirm the findings of Zuccarello et al. (2004), placing *Choreocolax, Harveyella* and *Leachiella* within the Rhodomelaceae. Furthermore, the phylogeny utilizing additional molecular markers provides strong support for a clade containing *Choreocolax, Harveyella* and *Leachiella* (Fig. 1), and supports the placing the Choreocolacaceae in synonymy with the Rhodomelaceae. Since their adoption by Feldmann and Feldmann (1958) the terms adelphoparasite and alloparasite have been used to describe parasites that infect hosts within their tribe/family, or in different tribes/families, respectively. The use of these terms has been questioned as molecular data have revealed that alloparasites, like adelphoparasites, infect close relatives rather than distantly related species (Zuccarello et al. 2004, Kurihara et al. 2010). To date, there is only one documented case of intrafamilial infections by red algal parasites that are supported by molecular systematics of host and parasite (Zuccarello et al. 2004). By placing the family Choreocolacaceae in synonymy with the Rhodomelaceae we are making steps to move away from a taxonomic definition for red algal parasites in favor of a biological distinction.

### A Biological Basis for Distinguishing Red Algal Parasites

With more than120 described parasites occurring across eight Florideophyceae orders (Salomaki and Lane 2014, Blouin and Lane 2015, Preuss et al. 2017, Preuss and Zuccarello 2018), red algae appear more able to transition from autotrophy to parasitic lifestyles than any other eukaryotic lineage. The data presented here place *Choreocolax polysiphoniae, Harveyella mirabilis*, and *Leachiella pacifica* firmly within the same family as their hosts, further supporting abandonment of the adelpho/allo-terms for differentiating two types of red algal parasites. However, differences between two groups of parasites remain, including their developmental patterns as they infect their hosts, and the origins of organelles (Table 2). We believe that it is time to abandon the terms adelpho- and alloparasite, which are based upon a taxonomic framework that is different today than it was when these terms were introduced. Instead we propose to distinguish red algal parasites by biological characteristics that can be easily tested using molecular techniques, the retention of a native plastid. Throughout the remainder of this manuscript we will use the term “archaeplastic” parasite to refer to those parasites that retain a native plastid (formerly most alloparasites), and “neoplastic” parasite to discuss those that hijack a host plastid rather than retain their own copy (formerly adelphoparasites).

**Table 2.** Distinguishing features between Archaeplastic and Neoplastic parasites.

|  | Archaeoplastic parasites | Neoplastic parasites |
| --- | --- | --- |
| Evolutionary origins | Monophyletic clades | Independent evolutionary events |
| Proliferation through host | Mitotically dividing rhizoidal filaments that directly fuse with host cells | Directly infect an individual host cell and subsequently spread from host cell to host cell via cell fusions |
| Site of nuclear DNA replication | Mitotically dividing parasite rhizoidal filaments | Host derived heterokaryon cells |
| Mitochondrion | Maintain complete and conserved native mitochondrion | Maintain complete and conserved native mitochondrion |
| Plastid | Retain a native plastid with reduced genetic capacity (loss of photosynthesis genes) | Utilize a host derived dedifferentiated plastid which is incorporated into parasite spores |

### Developmental differences

It had long been recognized that the association between parasitic red algae and their hosts was facilitated, at least in part, by the ability of red algae to form cell to cell fusions between adjacent, non-daughter cells, called secondary pit connections (Sturch 1899, 1926, Feldmann and Feldmann 1958). With modern microscopy, the structure and formation of secondary pit connections between parasites and their hosts was determined (Kugrens and West 1973, Goff and Cole 1976a, Wetherbee and Quirk 1982a, 1982b, Wetherbee et al. 1984). Furthermore, these cell to cell fusions were proposed to serve as the mechanism by which nutrients are transported from the host to parasite cells (Wetherbee and Quirk 1982b). With the use of epifluorescence microscopy, Goff went on to establish their importance in the infection process by demonstrating that red algal parasites are able to transfer their nuclei and organelles into host cells via secondary pit connections (Goff and Coleman 1984, 1985, 1987a, Goff and Zuccarello 1994).

In addition to recognizing the role of secondary pit connections in the infection process, sophisticated microscopy advanced our understanding of how parasitic red algae spread throughout their hosts and characterized the host responses. In a groundbreaking series of manuscripts, Goff described in great detail the biology of *Harveyella mirabilis* (Reinsch) F. Schmitz et Reinke, including its development, structure, and nutrient acquisition from its host *Odonthalia flocossa* (Esper) Falkenberg (Goff and Cole 1973, 1975, 1976a, 1976b, Goff 1976, 1979a, 1979b). These investigations provided a framework to more intimately understand the development and interactions of a range of red algal parasites.

Aside from the formation of secondary pit connections that initiate host infection, important differences in developmental patterns and their subsequent spread throughout the hosts have been observed between the two types of parasites (reviewed in Salomaki and Lane 2014, Freese and Lane 2017). First, archaeplastic parasites including *Choreocolax, Harveyella*, and *Holmsella*, spread throughout their hosts by producing mitotically dividing rhizoidal filaments that grow among host cells (Sturch 1899, 1926, Goff and Cole 1976a, Fredericq and Hommersand 1990). In contrast, the neoplastic parasites *Gracilariophila oryzoides, Gardneriella tuberifera*, and *Janczewskia gardneri*, infect their host cells directly and spread from cell to cell, rarely creating their own rhizoidal filaments (Goff and Coleman 1987a, Goff and Zuccarello 1994). Additionally, while neoplastic parasites appear to transform infected host cells and undergo nuclear replication solely within host cells (Goff and Coleman 1987a, Goff and Zuccarello 1994), archaeplastic parasites appear to only undergo DNA replication in their rhizoidal filaments and infected host cells maintain a 1:1 ratio of parasite nuclei to secondary pit connections (Goff and Coleman 1984, 1985, 1987b). These key developmental differences further lend support for a means of discussing parasitic red algae in a biological rather than a taxonomic context.

### Organellar Origins

The combination of microscopy with molecular tools challenged the dogma of how red algal parasites interact with their hosts. In a study investigating the origins of organelles from three neoplastic parasites, *Gracilariophila oryzoides, Gardneriella tuberifera’* and *Plocamiocolax pulvinata*, Goff and Coleman (1995) demonstrated that the parasites retain their own mitochondria. However all three had lost their native plastid, and instead incorporate a dedifferentiated host plastid when packaging their spores. Subsequent investigations into parasite mitochondria also implicated gene loss in the adelphoparasites *Gracilariophila oryzoides* and *Plocamiocolax pulvinata* (Hancock et al. 2010). In their study, *atp*8 and *sdh*C were described as pseudogenes in the *G. oryzoides* mitochondrion and *atp*8 was noted as missing completely from the *P. pulvinata* mitochondrion genome (Hancock et al. 2010). These losses were attributed to decreased selective pressures as the parasites increasingly relied on their hosts for ATP and other molecules essential to their survival. A more recent investigation spurred by those results, demonstrated that gene loss in the parasite mitochondria were the result of sequencing errors and/or downstream analysis (Salomaki and Lane 2016).

Red algal parasite plastids appear to be more dynamic and perhaps the more interesting story in red algal parasite evolution. Molecular investigations including deep genomic sequencing, have confirmed that all adelphoparasites investigated to date, *Faucheocolax attenuata, Gracilariophila oryzoides, Gardneriella tuberifera*, *Janczewskia gardneri*, and *Plocamiocolax pulvinata* do not retain a native plastid (Goff and Coleman 1995, Salomaki and Lane, unpublished). Previously, the plastid genome has been completely sequenced from the archaeplastic parasite *Choreocolax polysiphoniae* (Salomaki et al. 2015), and here we present the plastid genome of *Harveyella mirabilis*, and partial plastid genome sequences of *Leachiella pacifica*. Both completely sequenced plastids are greatly reduced in coding capacity compared to the plastids of free-living photosynthetic red algae, having lost all genes related to photosynthesis (Salomaki et al. 2015). Interestingly, genes involved in amino acid and fatty acid biosynthesis, iron-sulfur cluster synthesis, as well as transcription and translation are conserved (Salomaki et al. 2015). Aside from the loss of photosynthesis genes, the *H. mirabilis* plastid retains a high level of synteny with plastids of closely related free-living red algae, however the *C. polysiphoniae* plastid has experienced extensive genome reorganization (Figs. 3 & 4).

Developmental differences are notable, including the location of parasite DNA replication and mechanism of spreading throughout the host, but plastid origin appears to be central to differences observed in red algal parasite evolution. We hypothesize that the observed developmental differences between the traditionally termed adelpho- and alloparasites are intimately linked to plastid origin. Parasites that maintain a native plastid are, in essence, a complete red alga in their own right, while those that incorporate a host derived plastid are being pieced together with organelles from another organism they remain compatible with. By retaining their own plastid, the plastid remains under the same selective pressures as the rest of the parasite genome, and therefore, retains its ability to function for amino acid, fatty acid, protein, and isoprene biosynthesis – all necessary functions for survival. Those parasites that retain their own plastid are therefore capable of developing and functioning similar to other free-living red algae, with the exception of not being able to photosynthesize. Furthermore, the maintenance of a native plastid and the resulting self-reliance would enable those parasites to endure through evolutionary time, successfully adapting to new hosts and speciating. This hypothesis can be tested in future studies by investigating the organellar origins of parasites in the genus *Holmsella. Holmsella pachyderma* was previously placed in the Choreocolacaceae based upon its developmental patterns within its host, until 1990 when, along with *Holmsella australis*, it was transferred to the Gracilariaceae based upon morphological characteristics (Fredericq and Hommersand 1990). A subsequent phylogenetic analysis including *H. australis* and *H. pachyderma* found that the species formed a monophyletic clade within the Gracilariaceae (Zuccarello et al. 2004). To date no work has been completed on the plastid origin of members of the genus *Holmsella*, but if they do retain a native plastid, *Holmsella* would represent a second successful evolution of parasites retaining archaeplastic characteristics: a native plastid, mitotically dividing rhizoidal filaments as a means of vegetative growth throughout their hosts, and forming a monophyletic clade of parasite species. Further investigation of the taxonomic affiliation of parasite genera in families outside the Rhodomelaceae including *Gelidiocolax* and *Pterocladiophila* is warranted.

### Origin of Parasites

Two hypotheses have been proposed for the origin of red algal parasites. Setchell (1918) suggested that a mutation in a spore causes parasites to arise sympatrically, whereas Sturch (1926) postulated that parasitic red algae start out as epiphytes that become endophytes over time and increasingly rely on the host for nutrition. It seems plausible that Setchell’s origin hypothesis could explain the rise of the neoplastic parasites, which comprise the majority of known red algal parasite biodiversity. By evolving from their hosts, neoplastic parasites could more easily incorporate a copy of a genetically similar plastid as their own. The adaptation of incorporating a host plastid may provide a quick fix for what otherwise would have been a lethal mutation. The unique parasite nucleus and mitochondrion could continue spreading from cell to cell of its newly acquired host (and savior) via secondary pit connections, essentially transforming the host cells as has been described (Goff and Coleman 1985, Goff and Zuccarello 1994). Although providing a means of short-term survival for the newly transitioned parasite, co-opting a host plastid as their own would also establish these parasites on an irreversible path towards their own extinction. In order for a nonphotosynthetic parasite to survive it must retain compatibility for other plastid functions, including amino acid and fatty acid biosynthesis. Our current understanding suggests that the host-derived plastid is newly acquired during each new infection, therefore, the host plastid experiences one set of evolutionary pressures while the parasite nucleus evolves and accumulates mutations of its own. Eventually the parasite will inevitably lose the ability to communicate with the host plastid as the parasite and host increasingly become genetically distinct. This leaves the parasite with two possibilities, either find another host with a compatible plastid or go extinct. Alternatively, the success of an archaeplastic parasite may be explained by Sturch’s hypothesis. By evolving from a closely related epiphyte that is able to create secondary pit connections, the parasite may retain its own plastid and therefore increase its longevity and the opportunity to speciate as the parasite adapts to new hosts. Therefore, what were previously viewed as competing hypotheses to explain the evolution of red algal parasites, may each explain how different types of parasites arose.

### Conclusion

Following the earliest investigations of red algal parasites, two groups were recognized based on morphological distinctions and they were initially separated by taxonomic affiliation with their hosts (Setchell 1918, Feldmann and Feldmann 1958, Goff 1982). The use of molecular phylogenetics for evaluating evolutionary relationships has altered the taxonomic framework that was originally used for distinguishing red algal parasites. Distinct differences remain, however, and whether or not the parasite retains a native plastid or incorporates one from its host appears to be fundamental to parasite biology. We propose that the terminology for discriminating between red algal parasites should to reflect these biological differences, and identifying the plastid source has become substantially easier with modern sequencing technologies. Therefore, we recommend applying the term archaeplastic parasite to describe those that retain a native plastid throughout the infection cycle, and neoplastic parasite to those that have lost their native plastid and instead incorporate a host plastid. We predict that investigations of parasites that have traditionally been referred to as alloparasites in families other than the Rhodomelaceae, like *Holmsella* and *Pterocladiophila*, will also provide evidence of plastid retention and monophyletic clades of parasites.

## ACKNOWLEDGEMENTS

The authors thank Craig Schneider and Gary Saunders for conversation regarding taxonomy and names for distinguishing parasites. Jillian Freese is thanked for collecting specimens of *Harveyella mirabilis* and *Leachiella pacifica*, and Kristina Terpis for her support in the lab. Funding for this work was provided to by a Phycological Society of America Grant-in-Aid of Research to ES and by grant #1257472 from the National Science Foundation awarded to CL. This research is based in part upon work conducted using the Rhode Island Genomics and Sequencing Center, which is supported in part by the National Science Foundation (MRI Grant No. DBI-0215393 and EPSCoR Grant Nos. 0554548 & EPS-1004057), the US Department of Agriculture (Grant Nos. 2002-34438-12688 and 2003-34438-13111), and the University of Rhode Island.

## Abbreviations

ORF: = Open Reading Frame
BI: = Bayesian inference
MCMCMC: = Metropolis-coupled Markov chain Monte Carlo

